# Preserved Barrier Integrity and Altered Immune Profiles in Chronic Cannabis Users: Potential Roles of Δ9-Tetrahydrocannabinol

**DOI:** 10.64898/2026.07.02.736074

**Authors:** John E. McKinnon, Zejun Zhou, Amanda Wagner, Zhenwu Luo, Alicia Hartley, Zhuang Wan, Sylvia Fitting, Azizul Haque, Aimee McRae-Clark, Wei Jiang

## Abstract

Although cannabinoids such as delta-9-tetrahydrocannabinol (THC) are generally immunosuppressive in preclinical models, chronic cannabis use in humans is paradoxically associated with increased infection risk and systemic inflammation. In this study, we demonstrate that THC directly strengthens intestinal epithelial barrier function *in vitro* by increasing trans-epithelial electrical resistance in a concentration-dependent manner in Caco-2 monolayers. In a cross-sectional study of chronic cannabis users via smoking or snorting compared with non-using controls, plasma lipopolysaccharide (LPS), and microbial translocation-driven inflammatory cytokines (IL-23, MCP-1, IL-8) were significantly reduced, while some cytokines (IL-6, IL-1β, TNF-α, IL-10) remained unchanged. Concurrently, users exhibited elevated macrophage-derived chemokine (MDC) and homeostatic cytokines IL-15 and IL-21, markedly suppressed IL-7 and IL-4. Plasma IL-15 and MDC levels correlated with consumption intensity, and IL-23, IL-7, and IP-10 correlated with age of first use or during heaviest use. These findings suggest that habitual cannabis use may protect gut barrier integrity and reduce microbial translocation and associated inflammation, while simultaneously disrupting systemic immune homeostasis through selective cytokine dysregulation. This dual, dose-dependent immunomodulatory profile highlights the complex balance between potential benefits and risks in both recreational and therapeutic cannabis use.

## Introduction

Cannabis sativa is a chemically complex plant containing more than 560 identified compounds, including at least 104 cannabinoids. Among these, the most extensively studied are delta-9-tetrahydrocannabinol (THC) and cannabidiol (CBD). THC is the principal psychoactive constituent and exerts its effects primarily through activation of cannabinoid receptors, CB1, which is highly expressed in the central nervous system, and CB2, which is predominantly found in peripheral tissues [1]. These receptors, together with their endogenous lipid ligands (e.g., N-arachidonoylethanolamine [anandamide, AEA] and 2-arachidonylglycerol [2-AG]) and the enzymes responsible for their synthesis and degradation, constitute the endogenous cannabinoid system (ECS). In contrast, CBD is a non-psychoactive cannabinoid with distinct pharmacological properties [1].

Widespread cannabis use among adolescents and young adults is a significant public health concern, given its effects on brain regions involved in memory, learning, coordination, and sensory perception. Beyond its neurological effects, cannabis also modulates immune function. Preclinical and *in vitro* studies often show that cannabinoids such as THC suppress immune activation, inflammation, and macrophage functions [2, 3]. However, human data are less straightforward; chronic cannabis users frequently demonstrate increased susceptibility to infections and, in some cases, elevated systemic inflammatory markers, suggesting potential immune dysregulation [2]. The long-term impact of habitual cannabis exposure on immune homeostasis, particularly aspects linked to chronic low-grade inflammation, such as gut barrier integrity and microbial translocation, remains poorly defined.

In this study, we examined the effects of THC on intestinal barrier integrity and systemic immune status in chronic cannabis users. These findings provide new insight into how chronic cannabis use interfaces with gut barrier function and systemic immunity in humans.

## Materials and methods

### Study subjects

This study was approved by the Institutional Review Board (IRB) for Human Research at the Medical University of South Carolina. All participants were adults aged 19 and above and provided written consent. Participants in the cannabis group were current non-injection users with no documented prescription drug misuse or other illicit drug use, as verified by appropriate modules of the Mini-International Neuropsychiatric Interview and urine drug screening. Cannabis use patterns were assessed using the calendar-based, self-report Timeline Follow-Back (TLFB) method [4], which captured the frequency and quantity of cannabis consumption over the 90 days preceding the study visit. Participants reported whether they used cannabis (yes/no) and the number of joints, blunts, pipes, bowls, vaporizers, spliffs, edibles, or other methods consumed. When cannabis products were shared or only partially used, participants provided fractional counts. Demographics and clinical characteristics of this cohort are shown in Table 1.

### Plasma sample preparations

Plasma was isolated by centrifugation of EDTA-containing blood, aliquoted, and stored at -80°C until it was thawed for study.

### Plasma levels of LPS

Plasma samples were diluted to 10% with endotoxin-free water and heated to 80°C for 10 min to inactivate inhibitory proteins. LPS levels in plasma were then quantified using the Limulus amebocyte lysate QCL-1000 kit (Lonza, Walkersville, MD, USA) as described in our previous studies [5-7].

### Trans-epithelial electrical resistance (TEER)

Following our previous study [8], Caco-2 cells were cultured in 24-well plates and seeded at a density of 20,000 cells per insert onto 6.5-mm PET membranes with 0.4-µm pores (Corning, New York, USA). The culture medium was replaced every other day. Cells were treated for 60, 120, and 180 min with either control medium or THC (1, 10, and 30 µM). Following treatment, confluent monolayers were washed three times with pre-warmed PBS and allowed to equilibrate for 30 min at 37 °C. TEER was then measured using the Millicell Electrical Resistance System (ERS; Millipore, Billerica, MA, USA). All experiments were performed in triplicate.

### Plasma levels of proinflammatory and homeostatic cytokines and chemokines

Plasma concentrations of microbial translocation-associated proinflammatory cytokines and chemokines, including interleukin (IL)-1β, IL-8, IL-10, IL-23, IL-17A, tumor necrosis factor (TNF)-α, interferon gamma–induced protein 10 (IP-10), monocyte chemoattractant protein (MCP)-1, and macrophage-derived chemokine (MDC) were quantified alongside key homeostatic cytokines (IL-2, IL-4, IL-6, IL-7, IL-15, IFN-γ, and IL-21). All analyses were measured using the Human ProInflam Kit (Meso Scale Diagnostics, Rockville, MD), following the manufacturer’s instructions.

### Statistical analysis

Statistical analysis was performed by GraphPad Prism 6.0 (GraphPad, San Diego, USA) using the Student’s *t* tests, ANOVA, and Spearman correlation tests. P values of ≤ 0.05 were considered statistically significant.

## Results

### THC directly strengthens intestinal epithelial barrier function *in vitro*

Differentiated Caco-2 monolayers were treated with vehicle (0.1% ethanol), 1 µM, 10 µM, or 30 µM THC for 180 min. THC produced an apparent concentration-dependent increase in TEER (Figure 1A-1B). At 30 µM, TEER reached the highest level at 180 min treatment (p < 0.001 vs. vehicle, n = 6 inserts per condition, experiment repeated three times). No cytotoxicity was observed at any concentration tested (LDH assay < 5% release).

**Figure 1.**
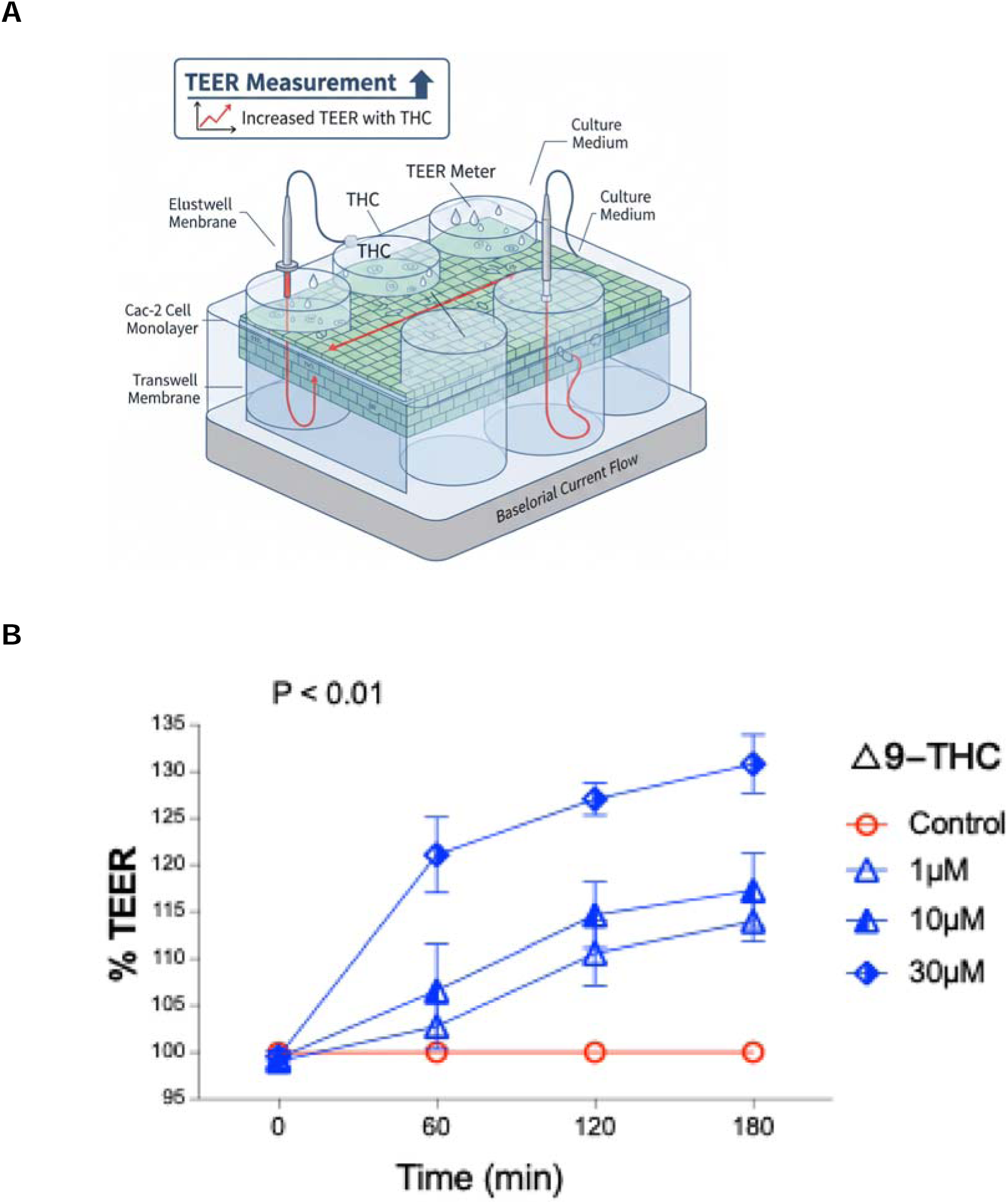
THC protects gut epithelial integrity *in vitro*. **(A)** Schematic of Caco-2 transwell system (Gemini). **(B)** Concentration–response curve of THC (1-30 µM) on TEER after 180 min treatment. Data are mean ± SEM from three independent experiments. p < 0.01 vs. vehicle. ANOVA.

### Chronic cannabis users display reduced systemic microbial translocation

To investigate the impact of THC on barrier integrity in humans, we evaluated plasma LPS, a robust marker of microbial translocation [9]. Notably, plasma LPS was significantly lower in chronic cannabis users than in non-using controls (P = 0.045) (Figure 2A-2B), which can be a consequence of cannabis-mediated barrier protection.

**Figure 2.**
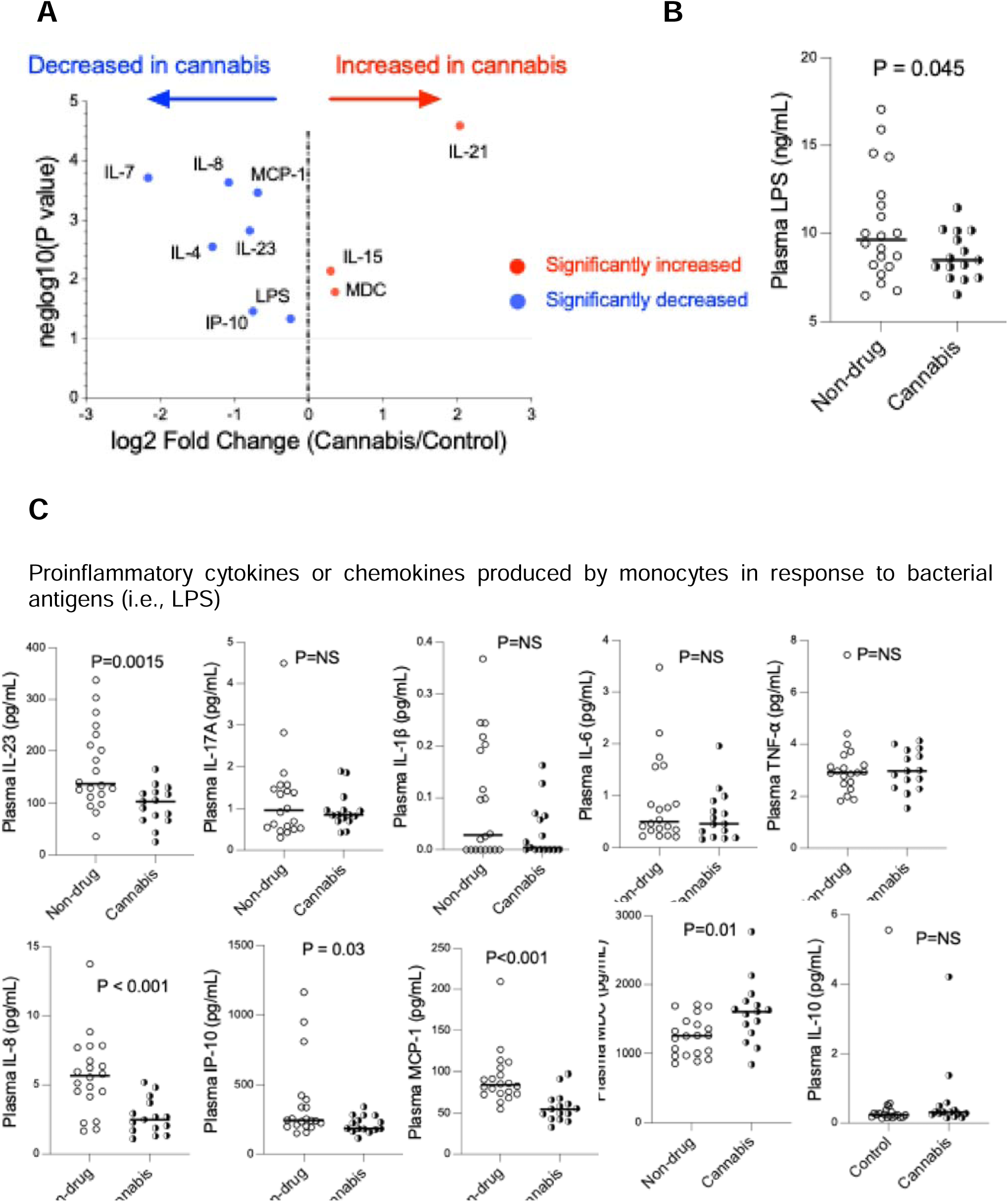
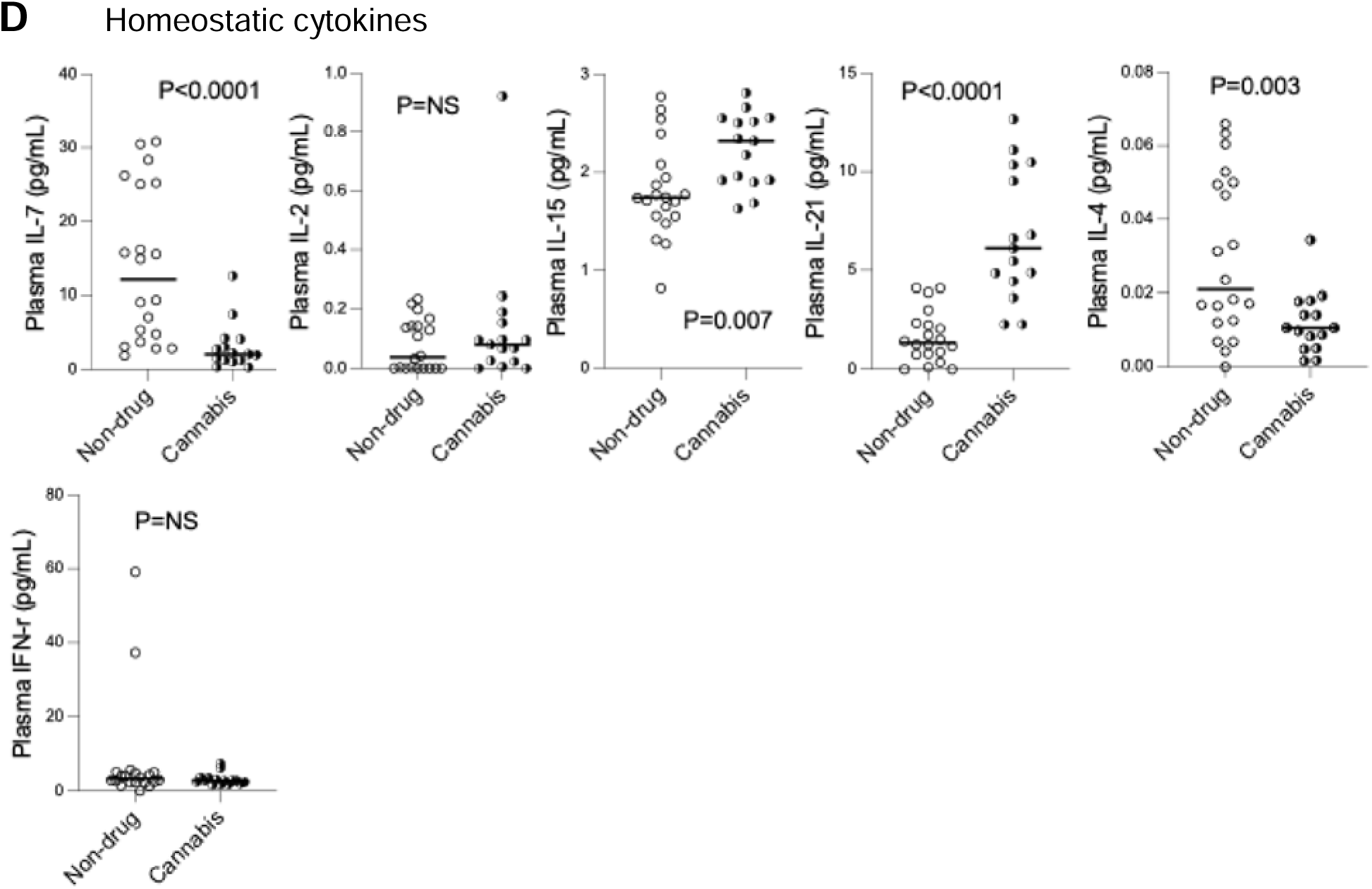
Decreased plasma levels of LPS and MT-related proinflammatory cytokines, increased chemokine MDC, and increased homeostatic cytokines IL-21 and IL-15 and decreased IL-7 and IL-4 in cannabis users versus controls. **(A)** Volcano plot displaying the log_2_ Fold Change (Cannabis/Control) on the x-axis against the log_10_ (P value) on the y-axis for various cytokines, chemokines, and LPS. Red dots indicate significantly increased markers in cannabis users (e.g., IL-21, IL-15, MDC), blue dots indicate significantly decreased markers (e.g., IL-7, IL-8, MCP-1, IL-23, IL-4, IL-18, IL-10, IP-10, LPS), and black dots are not significant. **(B)** Plasma LPS concentrations (ng/mL, median) in two study groups. **(C)** Plasma levels of proinflammatory cytokines and chemokines (pg/mL, median). Significant decreases in cannabis users are observed for IL-23, IL-8, IP-10, and MCP-1. MDC is significantly increased. IL-17A, IL-1β, IL-6, TNF-α, and IL-10 show no significant difference (P=NS). **(D)** Cannabis users exhibited decreased plasma homeostatic cytokines (pg/mL, median) IL-7 and IL-4, and increased IL-15 and IL-21. IL-2 levels are not significantly different. Two-tailed Student’s *t* tests.

### Selective suppression of microbial translocation-associated proinflammatory cytokines

Among the ten cytokines/chemokines quantified by MSD multiplex assay, four proinflammatory cytokines known to be induced by bacterial products were significantly reduced in cannabis users versus controls: IL-23 (P = 0.002), MCP-1/CCL2 (P < 0.001), IP-10 (P = 0.03), and IL-8/CXCL8 (P < 0.001) (Figure 2A and 2C). In contrast, LPS-induced classical proinflammatory cytokines IL-6, IL-1β, TNF-α, and anti-inflammatory IL-10 were not different between groups (all P > 0.05).

### Dysregulated homeostatic and myeloid-derived cytokines and chemokines

Next, we evaluated six key homeostatic cytokines, IL-7, IL-2, IL-15, IL-4, IFN-γ, and IL-21, which regulate the development of various T cell subsets, B cells, and myeloid cells. Cannabis users exhibited significantly higher levels of MDC/CCL22 (P = 0.01), IL-15 (P = 0.007), and IL-21 (P < 0.0001), but lower IL-7 (P < 0.0001) and IL-4 (P = 0.003) (Figure 2D). Plasma levels of IL-2 and IFN-γ remained unchanged (Figure 2D).

### Intensity of cannabis use correlates with IL-15, IP-10, IL-7, IL-23, and MDC levels

To investigate potential associations between cannabis use intensity and systemic immune parameters that differed between the two study groups, we conducted non-parametric Spearman rank correlation analyses between various cannabis use parameters and plasma levels of LPS and altered cytokines/chemokines (IL-4, IL-7, IL-8, IL-15, IL-21, IL-23, IP-10, MCP-1, and MDC). The overall correlation patterns are shown in the heatmap (Figure 3A), with significant associations (P < 0.05) indicated by asterisks. Positive correlations were observed between plasma IL-15 levels and THC quantity during heaviest use (r = 0.60, P = 0.03); between plasma MDC levels and both the current quantity per using day (including standardized measures) (r = 0.58, P = 0.025) and the quantity per using day during the first or heaviest use period (r = 0.54, P = 0.04) (Figure 3B). Significant negative correlations included plasma IP-10 levels and the age of first use (r = -0.65, P = 0.01), plasma IL-23 levels and age during the heaviest use period (r = -0.60, P = 0.02), and plasma IL-7 levels and age of first use (pattern) (r = -0.65, P = 0.01) (Figure 3C). No significant correlations were found with LPS or the other markers analyzed.

**Figure 3.**
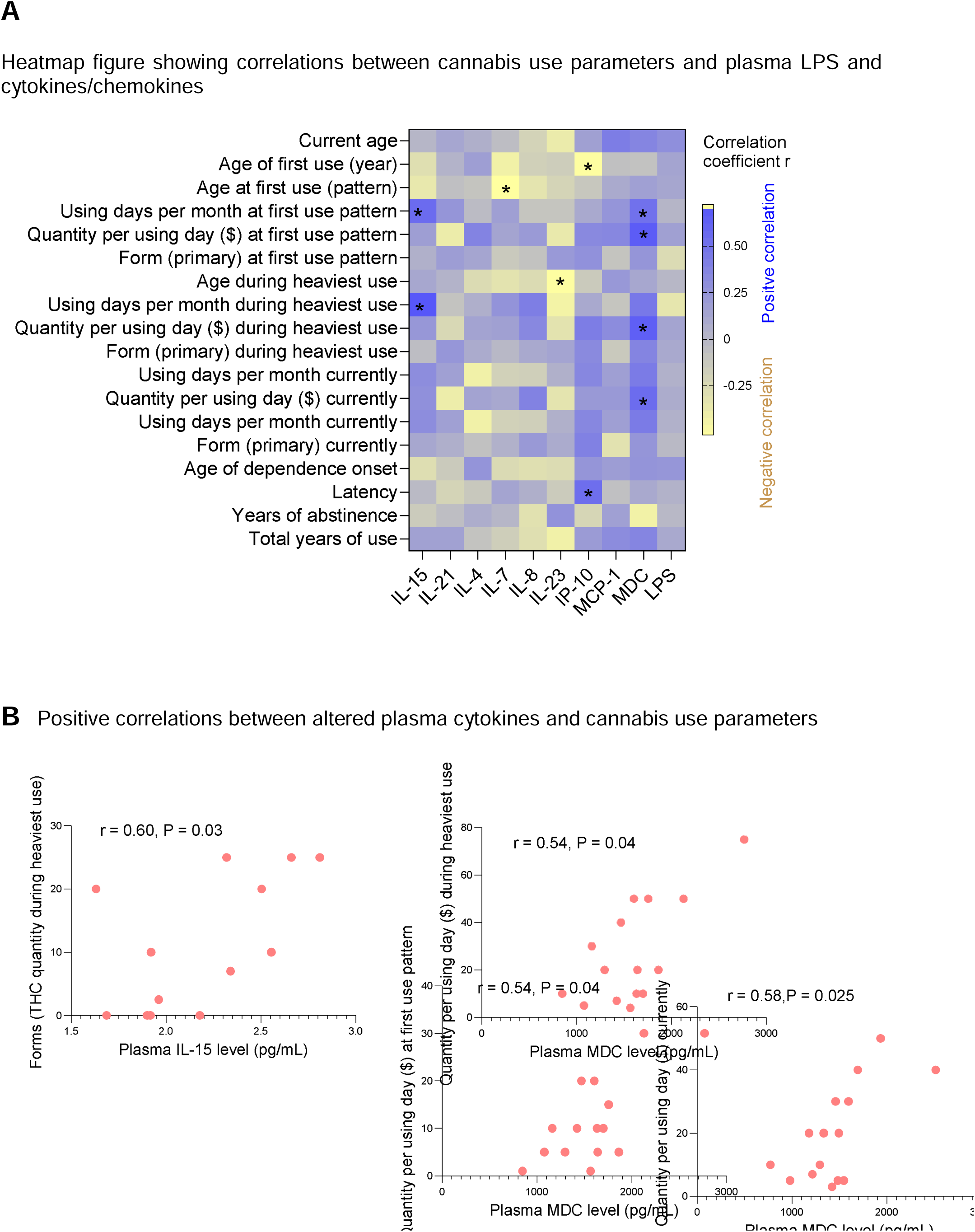
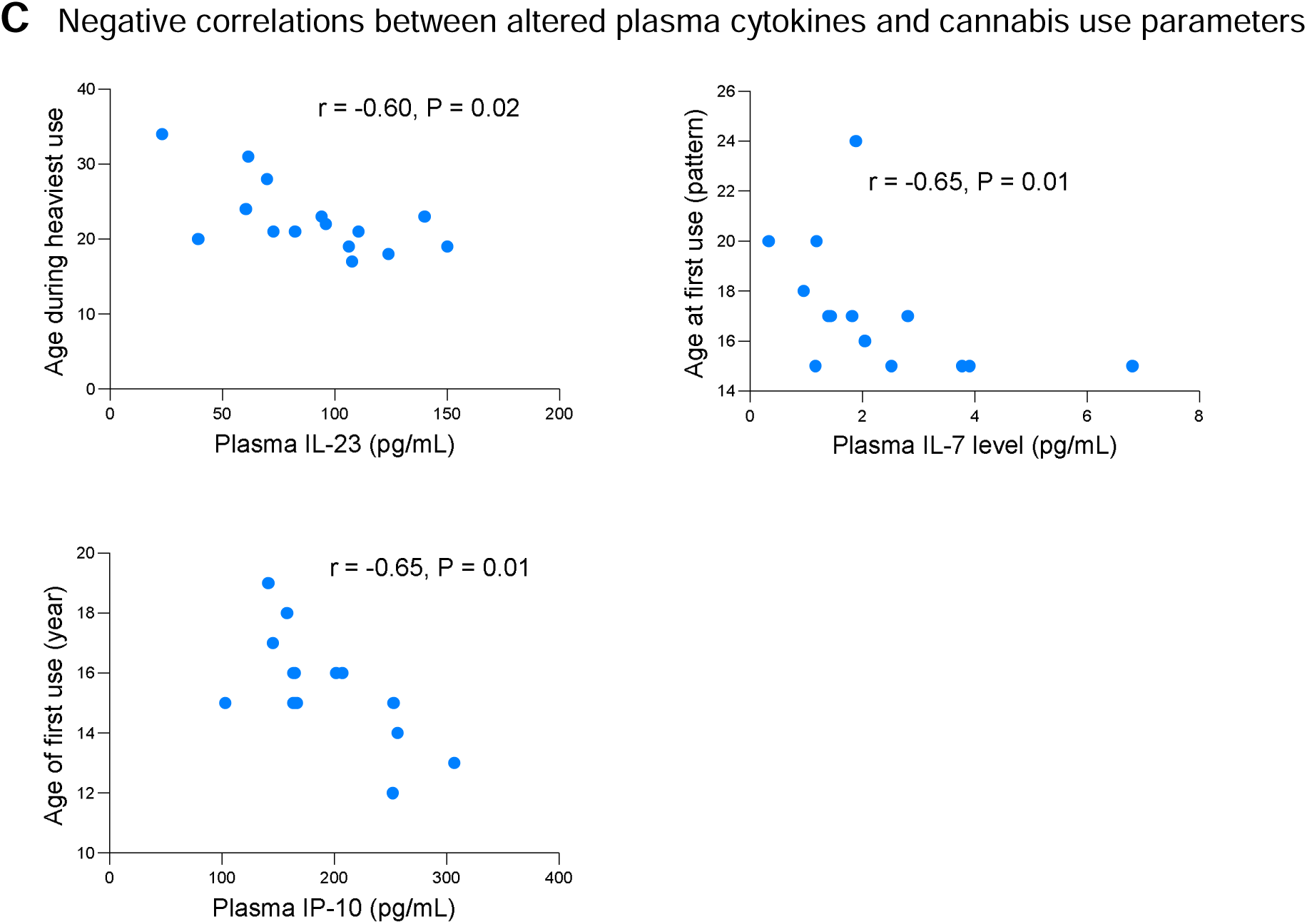
Correlations between indices of cannabis use and plasma levels of cytokines, chemokines, and LPS. **(A)** Heatmap showing Spearman correlation coefficients (r) between cannabis use parameters (rows) and plasma markers differed in the two study groups (columns). Blue indicates positive correlations, yellow indicates negative correlations, and only * indicates significant correlations (P < 0.05). **(B)** Positive correlations between altered plasma cytokines and cannabis use parameters, illustrated via scatter plots with trend lines, r values, and P values. **(C)** Negative correlations between altered plasma cytokines and cannabis use parameters, illustrated via scatter plots with trend lines, r values, and P values. Spearman correlation tests were used for all analyses.

## Discussion

This study provides the first integrated evidence, from *in vitro* barrier function to human *in vivo* biomarkers, that chronic cannabis use strengthens intestinal epithelial integrity and reduces microbial translocation-driven inflammation, while also dysregulating a homeostatic cytokine profile *in vivo*. The intensity of use or the younger age at first or heaviest use were correlated with higher plasma levels of IL-15, MDC, IL-23, IL-7, and IP-10, all of which can be produced by myeloid cells in response to microbial translocation.

In differentiated Caco-2 monolayers, THC treatment elicited a concentration-dependent enhancement of TEER, indicating strengthened barrier integrity without inducing cytotoxicity. This finding aligns with preclinical and *in vitro* data demonstrating that cannabinoids, including THC, can mitigate gut permeability induced by inflammatory stimuli through mechanisms that may involve CB1/CB2 receptor activation, modulation of tight junction proteins (e.g., claudins and occludin), or anti-inflammatory signaling pathways, such as NF-κB inhibition [10-12]. For instance, prior studies have shown that CB1 agonists reduce intestinal inflammation in murine colitis models by preserving epithelial homeostasis, suggesting a direct role for the endocannabinoid system (ECS) in gut barrier regulation [13].

Complementing these *in vitro* observations, our human cohort analysis revealed significantly lower plasma LPS levels in chronic cannabis users compared to non-users, a hallmark of diminished microbial translocation from the mucosal barrier. Elevated circulating LPS is implicated in driving low-grade systemic inflammation, which contributes to chronic inflammatory conditions like obesity, diabetes, and cardiovascular disease [14, 15]. The observed suppression of microbial translocation-associated proinflammatory mediators, IL-23, MCP-1, IP-10, and IL-8, further supports a lower inflammatory response to microbial products in cannabis users. These cytokines are typically induced by LPS via Toll-like receptor 4 (TLR4) signaling in monocytes, macrophages, and epithelial cells, promoting Th17 differentiation (IL-23), monocyte recruitment (MCP-1), and neutrophil chemotaxis (IL-8). Their selective reduction, without alterations in broader, classic proinflammatory (e.g., TNF-α, IL-1β, IL-6) or anti-inflammatory (IL-10) cytokines, suggests that chronic cannabis exposure may preferentially target microbial translocation-driven pathways rather than induce global immunosuppression. This pattern contrasts with some epidemiological reports of increased infection susceptibility in heavy cannabis users, which may stem from respiratory rather than systemic immune effects, or confounding factors like concurrent tobacco use. Moreover, IL-15, increased in users, is an inflammatory cytokine, whereas IL-4, decreased in users, is an anti-inflammatory cytokine that promotes M2 macrophage differentiation [16]; thus, chronic cannabis use is associated with a non-classical form of inflammation. Nonetheless, treatment of THC or CBD in human THP-1 macrophages suppressed IL-6, IL-10, and MCP-1, whereas IL-1β, IL-8, and TNF-α were less responsive [17]; the differences with our results may stem from *in vitro* versus *in vivo*, human primary samples versus a cell line, and acute versus chronic effects.

The observed pattern of decreased plasma levels of LPS and select proinflammatory mediators (IL-23, MCP-1, IP-10, and IL-8) without alterations in IL-6, TNF-α, IL-1β, IFN-γ, or IL-10 suggests a selective modulation of inflammatory responses, likely tied to specific upstream triggers or signaling branches or direct versus indirect effects rather than broad suppression. Consistently, THC administration to patients with multiple sclerosis did not change serum levels of IL-10, IL-12, IFN-γ, or C-reactive protein [18]. Below are outlined mechanisms based on established immunology, focusing on LPS as the key initiator (via TLR4 activation on monocytes, macrophages, and dendritic cells.

1. Reduced systemic low-grade persistent LPS exposure as the primary driver. LPS enters circulation primarily through compromised barriers. Elevated LPS translocation can result in chronic inflammatory conditions and complications (i.e., neuroinflammation, metabolic disease) [19], whereas a germ-free condition without LPS leads to immunodeficiency in animals [20]. Reduced LPS in chronic cannabis users lowers overall systemic TLR4 stimulation, but if the reduction is moderate, it may disproportionately affect "secondary" or amplification-phase mediators like chemokines (MCP-1, IP-10, IL-8) and IL-23, which require sustained or higher-threshold LPS signaling, while classic MT-associated cytokines (TNFα, IL-1β, IL-6) from rapid macrophage activation remain stable. IL-10, as a feedback regulator, might stay unchanged to prevent over-suppression. Notably, lower LPS limits the recruitment of additional immune cells (e.g., monocytes via MCP-1, T cells via IP-10, neutrophils via IL-8), thereby reducing feed-forward loops that amplify these mediators, without affecting core innate responses. Indeed, gut-targeted therapies for patients with metabolic syndromes reduce LPS and associated chemokines, but do not fully resolve systemic inflammation [21].
2. Differential engagement of TLR4 signaling branches (MyD88 vs. TRIF). LPS binds TLR4, triggering two adapter pathways: a) MyD88-dependent (rapid, plasma membrane-based) pathway activates NF-κB and MAPKs, leading to quick production of TNFα, IL-1β, IL-6, IL-8, MCP-1, and IL-10. This is the dominant pathway for early proinflammatory responses. b) TRIF-dependent (delayed, endosomal) pathway activates IRF3/7 for type I interferons (IFN-β/α) and IFN-inducible genes like IP-10; also contributes to late NF-κB. IL-23 production often requires cooperation between MyD88 and TRIF for optimal induction in dendritic cells. If the TRIF pathway is preferentially dampened (e.g., by endocytosis inhibitors, specific adapters like TRAM, or conditions favoring MyD88 bias), it would reduce IP-10 (strongly IFN-dependent) and IL-23 (requires TRIF for full expression), while MyD88-driven cytokines (TNFα, IL-1β, IL-6, IL-10) remain intact. For MCP-1 and IL-8 (mostly MyD88), their decrease might stem from reduced TRIF-mediated amplification or lower overall LPS triggering thresholds. This branching explains why some LPS variants or modulators (e.g., low-acyl LPS forms) bias toward one pathway, leading to skewed cytokine profiles. In viral-bacterial co-infections or targeted therapies (e.g., TRIF inhibitors), TRIF suppression limits IFN-related mediators without abolishing core inflammation.
3. Cell-type-specific responses and feedback regulation. Cytokines originate from distinct cells, for example, macrophages/monocytes are the primary source of TNFα, IL-1β, IL-6, IL-10 (rapid LPS response); dendritic cells are key for IL-23 (Th17 promotion); epithelial/endothelial cells produce chemokines like IL-8 (neutrophil attractant), MCP-1 (monocyte attractant), and IP-10 (T cell attractant) in response to localized LPS or secondary signals [19]. Reduced LPS might preferentially limit activation of non-myeloid cells (e.g., gut epithelium), decreasing chemokine output, while myeloid cells maintain baseline cytokine production. IL-10, induced for homeostasis, remains stable to counterbalance. Reduced cell migration (due to lower chemokines) also prevents amplification cascades that boost IL-23 and chemokines without affecting primary cytokines. Notably, in resolving inflammation (e.g., post-antibiotic treatment in burns or sepsis), LPS clearance improves, but macrophage activation persists [22].
4. Potential contributing factors: kinetics and dose effects. Myeloid cell responses to LPS are highly dependent on exposure conditions, including low versus high dose, acute (single) versus chronic stimulation, and structural variations among bacterial LPS species [23]. Notably, low-dose LPS preferentially engages TRIF/IFN signaling pathways; thus, a reduction in LPS may initially favor TRIF activation while sparing higher-threshold MyD88-dependent cytokine responses. Negative Regulators may also play a role, such as upregulation of miRNAs (e.g., miR-146b, IL-10-dependent) or SOCS proteins, which could selectively repress chemokine genes without affecting others.

Cannabis users displayed dysregulated profiles of homeostatic and myeloid-derived cytokines/chemokines, including elevated MDC, IL-15, and IL-21, alongside decreased IL-7 and IL-4. MDC, produced by macrophages and dendritic cells, attracts Th2 cells and regulatory T cells, potentially fostering an anti-inflammatory milieu that aligns with reduced microbial translocation. Elevated IL-15 and IL-21, which support T cell and NK cell survival/proliferation, could indicate compensatory mechanisms to maintain immune vigilance amid suppressed proinflammatory signaling. Conversely, lower IL-7 (critical for lymphocyte homeostasis) and IL-4 (Th2 cytokine) might reflect impaired lymphoid maintenance or skewed polarization, possibly contributing to long-term immune dysregulation. Adolescent exposure occurs during a critical window when both the endocannabinoid system (ECS) and the immune system are actively maturing. Early disruption of the ECS during this developmental phase may permanently alter or "imprint" baseline immune tone and myeloid cell responsiveness, offering an explanation for why age of first use correlates with adult cytokine phenotypes. These alterations echo preclinical evidence where THC modulates cytokine networks via ECS interference with transcription factors like STATs or AP-1 [24].

Correlation analyses further linked cannabis use parameters to these immune shifts, revealing positive associations between initial use form/quantity metrics and IL-15/MDC levels, and negative correlations with age-related parameters and IP-10/IL-23/IL-7. Notably, earlier age of onset correlated with lower IL-7 and IP-10 (CXCL10, a T cell chemoattractant), suggesting that adolescent exposure, when the ECS and immune system are maturing, may imprint lasting homeostatic imbalances. Heavier use intensity positively correlated with MDC, implying dose-dependent effects on myeloid chemokine production. The absence of LPS correlations may indicate that barrier-protective effects saturate at moderate use levels and the chronic stage, while cytokine perturbations scale with exposure. These findings extend prior human studies, which have variably reported elevated C-reactive protein or altered T cell subsets in cannabis users, by highlighting gut-immune axis involvement [18].

The finding that chronic cannabis use enhances intestinal barrier integrity and reduces microbial translocation-driven inflammation has several clinically relevant implications. First, cannabinoid-based therapies may benefit conditions marked by gut-barrier dysfunction, including inflammatory bowel disease (IBD), irritable bowel syndrome (IBS), metabolic dysfunction-associated steatotic liver disease (MASLD), graft-versus-host disease (GVHD), obesity, type 2 diabetes, HIV, and chemotherapy-induced mucositis. The concentration-dependent barrier protection observed with THC *in vitro*, together with reduced plasma LPS in users, supports testing low-to-moderate THC or selective CB1/CB2 agonists in these settings, complementing ongoing trials in Crohn’s disease and ulcerative colitis. Second, in HIV and other chronic infections, the lower LPS and MT-related cytokines seen in cannabis users suggest a potential reduction in residual inflammation. However, concurrent reductions in IL-7 and IL-4 may impair lymphocyte homeostasis and vaccine responsiveness, requiring careful clinical counseling. Third, these data challenge simplified public health messages portraying cannabis as uniformly immunosuppressive; moderate-to-heavy use may simultaneously reduce systemic inflammation through improved gut-barrier function. Dose-, duration-, and route-specific guidance, favoring non-combustion delivery, may more accurately reflect the actual risk-benefit profile. Fourth, plasma MDC levels emerge as promising biomarkers of cannabinoid exposure intensity and could guide individualized dosing in future therapeutic trials. Finally, the decline in IL-7 warrants caution in adolescents, older adults, and immunocompromised patients, where long-term heavy use may exacerbate immunosenescence or susceptibility to infection.

This study has several limitations. The cross-sectional design precludes causality; sample sizes were modest (n = 15 users, n = 20 controls), potentially limiting power to detect subtle correlations. Cannabis use was self-reported via TLFB, which, while validated, may introduce recall bias, and we did not quantify CBD versus THC ratios or confirm chronicity via hair analysis. The cohort excluded polysubstance users, enhancing specificity but limiting generalizability. *In vitro* experiments used THC alone, not accounting for entourage effects from other cannabinoids. Future research should employ interventional trials with controlled THC dosing and frequency to validate barrier-protective effects in humans.

In summary, these findings indicate cannabis as a context-dependent immunomodulator that can confer barrier-protective, anti-inflammatory effects while posing risks of selective lymphoid dysregulation, underscoring the need for nuanced, evidence-based clinical integration. The net clinical effect likely depends on dose, route, and duration.

## Acknowledgements

This work was supported by the National Institute on Drug Abuse grant R01DA059854 (WJ), R01DA059538 (WJ), K24DA038240 (AM), and R01DA055523 (SF, WJ), and 1R21NS118393-01 (AH). This work was also supported in part by the 2026 Brain & Behavior Research Foundation (BBRF) Distinguished Investigator Award (34152, Jiang).

## Author contributions

Z.Z., A.W., and Z.L., recruited study participants and performed experiments. J.E.M and W.J. wrote the manuscript. Z.W. analyzed data. S.F., A.H., A.M.C., and W.J. revised the manuscript.

## Data availability statement

Data will be deposited during manuscript submission.

